# Could the recent zika epidemic have been predicted?

**DOI:** 10.1101/139253

**Authors:** Ángel G. Muñoz, Madeleine C. Thomson, Anna M. Stewart-Ibarra, Gabriel A. Vecchi, Xandre Chourio, Patricia Nájera, Zelda Moran, Xiaosong Yang

## Abstract

Given knowledge at the time, the recent 2015-2016 zika virus (ZIKV) epidemic probably could not have been predicted. Without the prior knowledge of ZIKV being already present in South America, and given the lack of understanding of key epidemiologic processes and long-term records of ZIKV cases in the continent, the best related prediction was for potential risk of an *Aedes*-borne disease epidemic. Here we use a recently published two-vector capacity model to assess the predictability of the conditions conducive to epidemics of diseases like zika, chikungunya or dengue, transmitted by the independent or concurrent presence of *Aedes aegypti* and *Aedes albopictus*. We compare the potential risk of transmission forcing the model with the observed climate and with state-of-the-art operational forecasts from the North American Multi Model Ensemble (NMME), finding that the predictive skill of this new seasonal forecast system is highest for multiple countries in Latin America and the Caribbean during the December-February and March-May seasons, and slightly lower –but still of potential use to decision-makers– for the rest of the year. In particular, we find that above-normal *suitable conditions* for the occurrence of the zika epidemic at the beginning of 2015 could have been successfully predicted for several zika hotspots, and in particular for Northeast Brazil: the heart of the epidemic. Nonetheless, the initiation and spread of an epidemic depends on the effect of multiple factors beyond climate conditions, and thus this type of approach must be considered as a guide and not as a formal predictive tool of vector-borne epidemics.

## 1 Introduction

Zika virus (ZIKV, family *Flaviviridae*, genus *flavivirus*) disease, a viral illness transmitted primarily by the *Aedes aegypti* and *Aedes albopictus* mosquitoes (1). ZIKV has recently emerged as a major epidemic in Latin America and the Caribbean, with 738,783 suspected and confirmed cases reported to date (2). Prior studies from Yapp Island suggest that the majority of ZIKV infections are asymptomatic or result in mild disease (3), and initial studies from Latin America suggest that the ZIKV infections are less severe and less febrile than chikungunya (CHIKV) or dengue (DENV) infections (4). The spread of ZIKV has been accompanied by severe neurological complications, including children born with microcephaly (5,6) and people with Guillain-Barré syndrome (7,8).

In a previous study (9), our team analyzed the potential contribution of climate signals acting at different timescales in creating the environmental scenario for the current ZIKV epidemic. We found that suitable climate conditions were present, due to the co-occurrence of anomalously high temperatures and persistent below-normal rainfall in several regions of South America, especially in Brazil, the heart of the epidemic.

These suitable conditions are not only favorable for ZIKV, but in general enhance the probability of both *Aedes sp.* reproduction and viral replication. Due to the fact that ZIKV, DENV and CHIKV share the same mosquito vectors and seem to have similar temperature dependence for their extrinsic incubation periods (10), there are advantages in considering the overall eco-epidemiological conditions for the potential risk of transmission of *Aedes*-borne arboviruses rather than focusing on the risk of transmission of only one disease. The effect of rainfall on *Aedes sp*. is more complex than temperature (e.g., (11,12)), because *Aedes* vectors breed in domestic water containers which are more abundant during droughts and water shortages (13). Their presence is also known to increase following unusually high rainfall when peri-domestic breeding sites (discarded containers, flower pots, tires, etc.) are filled with water.

The study of the different environment-virus-vector-human interactions in this field is normally performed using a diversity of mathematical models, most of them based on the Ross-McDonald model (14) or its generalizations. These models are commonly referred to as compartmental models, normally stratifying the human population in susceptible (S), infected (I) and recovered (R) individuals, which explains why they are also known with the alternative name of SIR models. A set of coupled differential equations is used to describe the evolution of each compartment (15,16). These models vary in complexity, and tend to be classified as homogeneous or heterogeneous models; for details, see for example (17).

Although these models are most frequently used to diagnose past or present epidemics, they can also be used in predictive mode, even at seasonal scale (see Thomson et al. (18) and references therein). Predicting environmental suitability conditions presents a complex problem, but it is indeed less complex than predicting the occurrence and transmission of the diseases in human populations. The complexity resides in the non-linear interactions between the different components of the coupled system in consideration, in which the effects of population immunity and susceptibility, or different possible immunological interactions between the diseases (e.g., co-infections of DENV and ZIKV) are still not well understood.

In this paper, we develop a new seasonal forecast system for suitable climate conditions conducive to enhanced transmission risk of *Ae. aegypti*- and *Ae. albopictus*-borne diseases. We use a two-vector model and state-of-the-art climate forecasts to assess its predictive skill, and we discuss the implications for Latin America and the Caribbean. For brevity, in the following pages we will use “potential risk of transmission” to refer to potential transmission associated with climate conditions suitable for transmission of the diseases mentioned earlier in this paper. Data and general methods are presented in Section 2, the vectorial capacity model is discussed in Section 3, the skill assessment for different seasons of the year is analyzed in Section 4, and the concluding remarks are presented in Section 5.

## 2 Data and Methods

The domain of study includes Latin America and the Caribbean, and is defined by the boundaries 120°W-25°W and 60°S-32°N.

The observed monthly temperature and rainfall fields for the period 1950-2015 were obtained from the University of East Anglia Climate Research Unit product version 3.24 (CRUv3.24; (19)), available at a horizontal resolution of 0.5 degrees. These datasets were selected to be consistent with our previous study on a similar topic (9). Tests indicated that the results are consistent with other large scale gridded climate datasets, such as the Climate Anomaly Monitoring System (CAMS, Global Historical Climatology Network version 2; see (20) for details) used in (21).

State-of-the-art temperature and rainfall forecasts at monthly timescales were obtained from the North American Multi-Model Ensemble project (NMME; (22)), at a common horizontal resolution of 1 degree. The total of 116 members available was used for the hindcast1 period of 1982-2010, but only 104 members were used for the December-February 2014-15 forecast, since those were the ones available for that year (no members from the NCAR-CESM^1^ and NASA-GMAO models). Hindcasts and forecast correspond to the month prior to the target season; for example, for the December-February season, the hindcast and forecast of November was used.

The vector model used in this work was recently developed by Caminade et al. (21). For the sake of organization, the vectorial capacity model equations are presented in the next section. The model requires climate information, and thus the observations and NMME forecasts mentioned above were used, the first one for diagnostics and the second one for the prognostic set up. The model was coded and executed in Matlab at a monthly timescale for a total of 792 months when forcing it with observed data, and 348 months per member when using the NMME hindcasts; each member was run independently before computing the ensemble and seasonal average. The vectorial capacity model output, forced with both climate observations and hindcasts, is available online in the Latin American Observatory’s Datoteca (23–26): http://datoteca.ole2.org/maproom/Sala_de_Salud-Clima/ContexHist-Map-1/index.html.es.

When analyzing the model forced with observations, standardization was performed with respect to the 1950-2015 period. Anomalies are defined as the value of the variable being analyzed minus its 1950-2015 average. To analyze inter-annual variability, a 12-month running average was computed. A linear detrending was used.

Skill was assessed using both Kendall’s τ and the 2AFC score (27), computed using the International Research Institute for Climate and Society (IRI) Climate Predictability Tool, CPT (28), version 15.4.7. Kendall’s τ is a non-parametric rank correlation coefficient used here to measure the overall association between observations and model output. The 2AFC score indicates the probability of correctly discriminating an observation in a higher category from one in a lower (e.g., an “above-normal” observation from a “normal” observation) given the forecasts expressed in deterministic form (i.e., the actual model values, and not the probabilities associated with them). The following four seasons were considered: December-February, DJF, March-May, MAM, June-August, JJA, and September-November, SON. A cross-validation window of five years was used, for the 1982-2010 period. For each iteration, five years were left out and the remainder years were used to build the statistical model, forecasting the middle year of the five-year window. This window is shifted one year into the future for the next iteration, and so on. The skill reported is the average of the metric computed for each iteration, and it was assessed after magnitude and spatial biases were corrected using a simple Model Output Statistics approach involving a Principal Component Regression (PCR; (29,30)), an option available in the CPT software. For further details see (29).

Maps showing the 2AFC score computed using this methodology were produced for each of the seasons considered. Categories for above normal, normal and below normal were identified in the vector model output using the typical 33.33% and 66.66% thresholds in the corresponding probability density function. Forecast probabilities for each category were computed using the PCR model built with the CPT package.

## 3 Two-vector ento-epidemiological model

Both *Ae. aegypti* and *Ae. albopictus* are considered the most important vectors in Latin America and the Caribbean for the transmission of ZIKV, CHIKV and DENV (e.g., (31–33)) These vectors are known to have different susceptibilities to these diseases, as well as different feeding characteristics (21). While *Ae. aegypti* and *Ae. albopictus* are considered a domestic mosquito and peri-domestic mosquito respectively, it is in principle possible to find them co-existing in the same place (34,35), something that is expected to be even more common in the near future due to global warming (31,36). Hence, we consider that an actionable seasonal forecast system like the one in consideration should involve at least these two species for Latin America and the Caribbean. This section presents the model equations used by the prediction system.

As it has been shown by other authors (21,37) the equations for the dynamics of a two-vector-one-host capacity model, a generalization of the well-known Ross-McDonald model (14), are

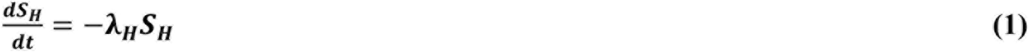

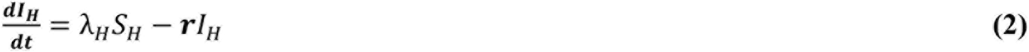

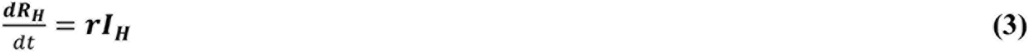

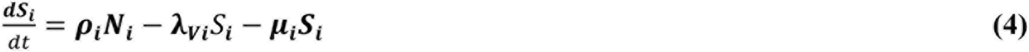

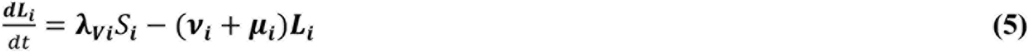

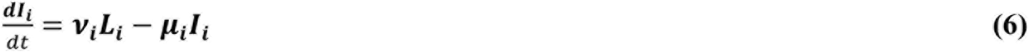

where ***S*_*H*_, *I*_*H*_** and ***R*_*H*_** are the number of susceptible, infectious and recovered hosts, respectively, associated with the *Aedes*-borne disease of interest. ***S*_*i*_, *L*_*i*_** and ***I*_*i*_** are the number of susceptible, latent and infectious vectors of kind ***i=1,2*** (*Ae. aegypti* and *Ae. albopictus*, respectively). In addition,

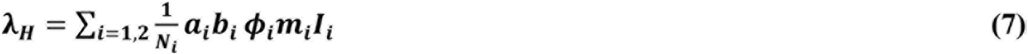

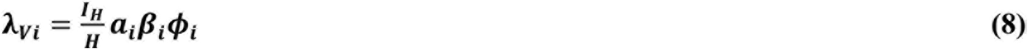

and ***a*_*i*_** is the daily biting rate (a function of temperature), ***b*_*i*_** is the vector-to-host transmission probability, ***ϕ_i_*** quantifies the vector’s preference for humans, ***m*_*i*_** is the vector-to-host ratio (a function of both temperature and rainfall; see (21) for details), *β*_*i*_ is the host-to-vector transmission probability, ***r*** is the daily recovery rate, and ***v*_*i*_** and ***μ_i_*** are the inverse of the extrinsic incubation period of the virus in days and the mortality rate^2^, respectively, both a function of temperature. As in (21), the vector-to-host-ratio ***m*_*i*_** is defined in terms of the probability of occurrence of the vectors (multiplied by 1000), which was obtained in (34) using maximum and minimum annual rainfall to account for the presence of water-filled containers, and other environmental variables involving temperature and urbanization; for details see the Materials and Methods section in (34). ***H*** and ***N*_*i*_** are the total number hosts and the total number of the *i-th* kind of vector, respectively.

This is a 5-compartment model: infectious human host, latent *Ae. aegypti* vectors, latent *Ae. albopictus* vectors, infectious *Ae. aegypti* vectors and infectious *Ae. albopictus* vectors. If **Δ** and **Λ** are the new infectious rate appearing in a compartment and the rate at which individuals leave said compartment, respectively, then

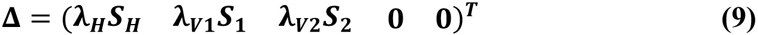

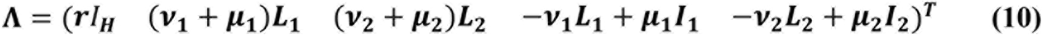

The basic reproduction number ***R*_0_** is the dominant eigen-value of the next-generation matrix (21)

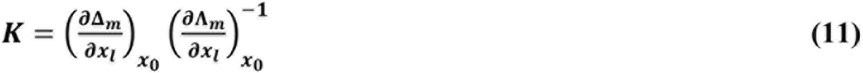

for ***m,l=1..5*** identifying the different compartments, ***x*** being a vector with the number of individuals in each compartment, and ***x*_0_** denoting the disease-free equilibrium state. The only non-zero elements ***K*_*ml*_** (new infections in compartment m produced by infectious individuals in compartment ***K*** are

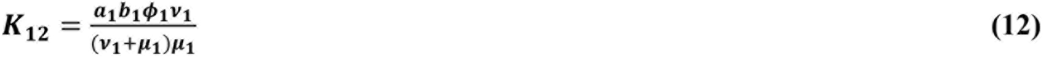

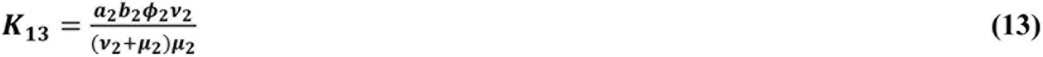

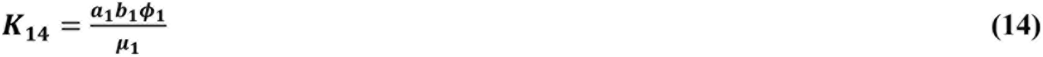

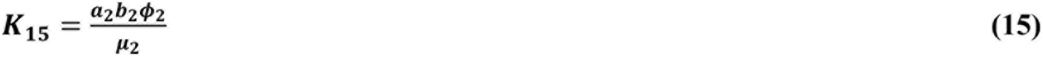

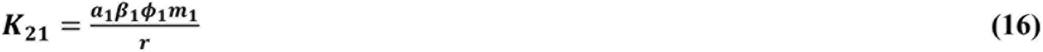

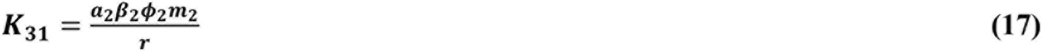

***R*_0_** is the largest eigen-value solution of the eigen-value problem **|*K* − *R*_0_*I*| = 0**:

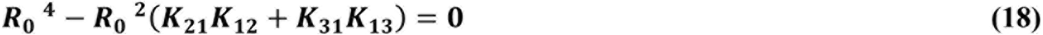

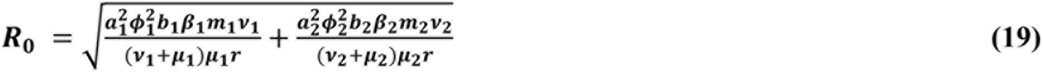

where, as the indices suggest, the first term in the square root corresponds to *Ae. aegypti* and the second one to *Ae. albopictus*. As in (21), we set ***R*_0_ = 0** in all locations and times for which the total monthly rainfall has not been at least 80 mm during a minimum of five months, a condition for stable transmission.

This model has been reported (21) to reproduce well the observed basic reproduction number obtained when using the relatively short record of ZIKV cases available in Latin America. Because of this, we have chosen the same values of the parameters and functional dependence on temperature and rainfall that was used in a prior study (21).

The basic reproduction number can be understood as the expected number of new cases generated by a single (typical) infection in a completely susceptible population. It is a dimensionless number that can be associated with the potential risk of transmission of the disease (in the sense of the suitable conditions for transmission, as discussed at the Introduction), considering only basic environmental, entomological and epidemiological information. Only values of ***R*_0_**>**1** are related to spreading the infection in the population, and thus we focus on that range of values of *R*_0_ in the present study.

The temperature dependence of certain parameters in the model (for example, the mortality rate ***μ_i_***; see Figure 1) strongly controls the spatial and temporal distribution of ***R*_0_**. Most of Latin America and the Caribbean typically exhibit high values of the two-vector basic reproduction number (Figure 2). The potential risk of transmission of *Aedes*-borne diseases is higher for the northern half of South America, especially in Brazil, most of Colombia, Venezuela, Guyana, Suriname and the French Guyana, coastal Ecuador and the Ecuadorian and Peruvian Amazon. Central America and the Caribbean, although to a lesser degree, also exhibit high values of ***R*_0_** Furthermore, with the increasing occurrence of high-temperature records, the frontier is extending farther into southern South America, in countries like Uruguay, which reported the first cases of autochthonous dengue fever in 2016 (38). Nonetheless, places that are too hot decrease the life expectancy of the vectors (roughly speaking, with temperatures above 40°C, see Figure 1), and thus some regions in the future could start seeing a relative decrease in vector number if temperatures keep increasing.

**Figure 1.**
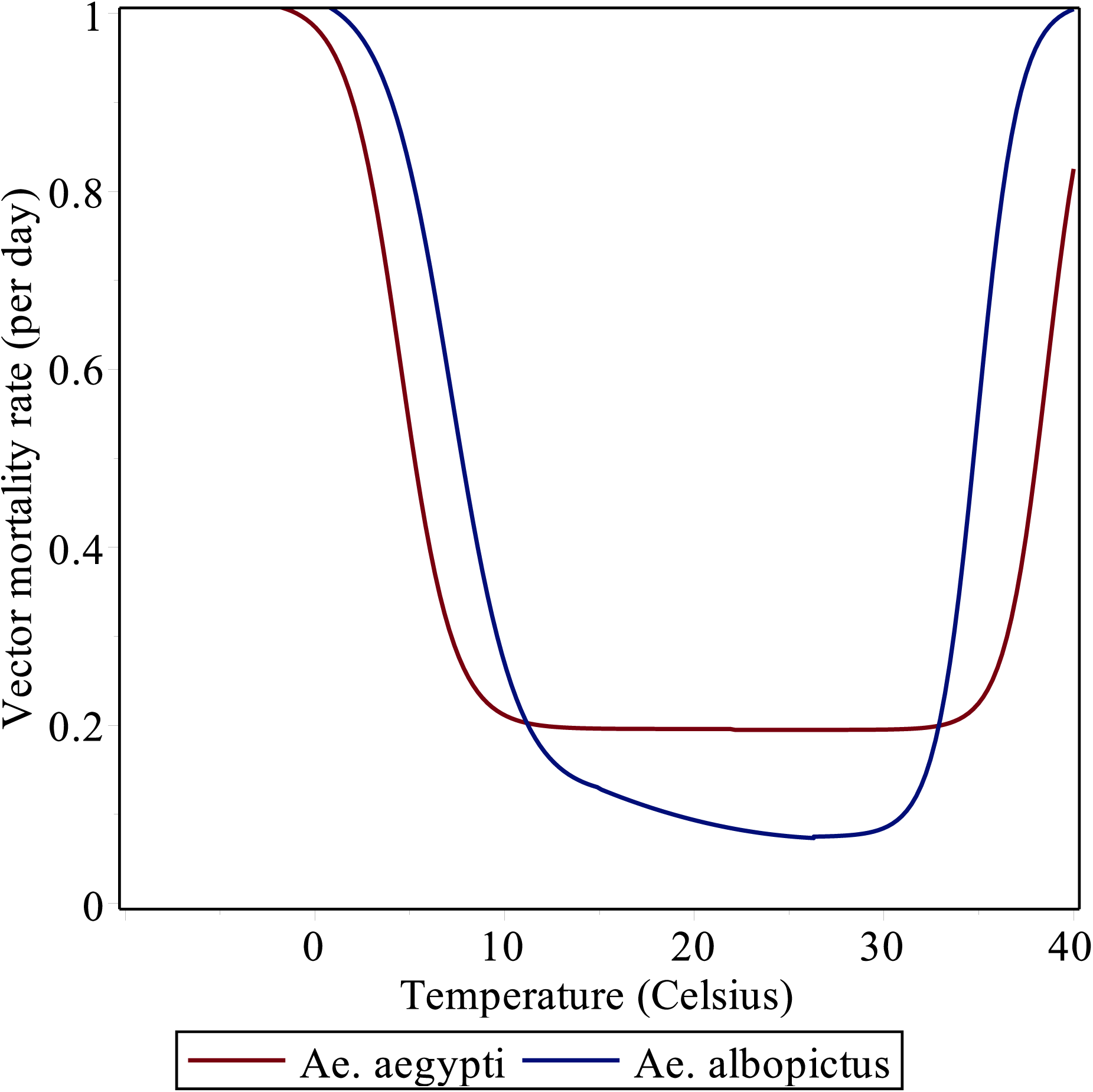
Daily vector mortality rate as a function of mean temperature (in Celsius).

**Figure 2.**
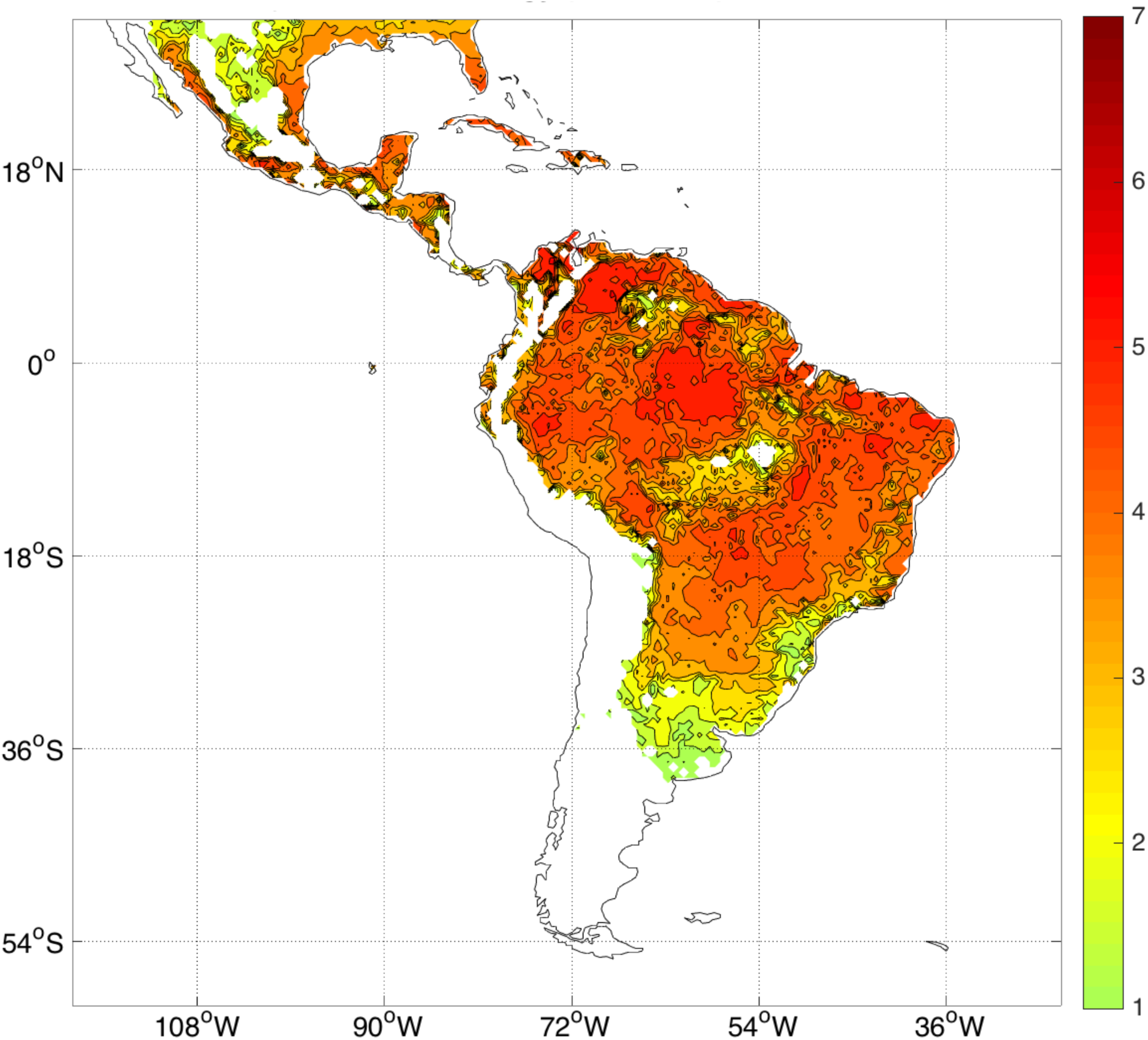
Observed climatology of ***R*_0_** considering all months in the period 1982-2010. Only ***R*_0_** > 1 values are plotted. There is no data over the oceans.

An analysis of the evolution of the suitable conditions for transmission during 2013-2015 (Figure 3) complements the study on the associated temperature and rainfall anomalies performed previously (9). Standardized positive ***R*_0_** anomalies covering regions of northern South America and northern Brazil in 2013 became dominant in almost everywhere in northern half of South America, Central America and the Caribbean in 2015, with values exceeding one standard deviation in zones of Brazilian Amazon, the northern Peruvian coast, all of coastal Ecuador, most of northern Colombia and western Venezuela. Standardized anomalies of around two standard deviations occurred in the heart of the Brazilian Amazon.

**Figure 3.**
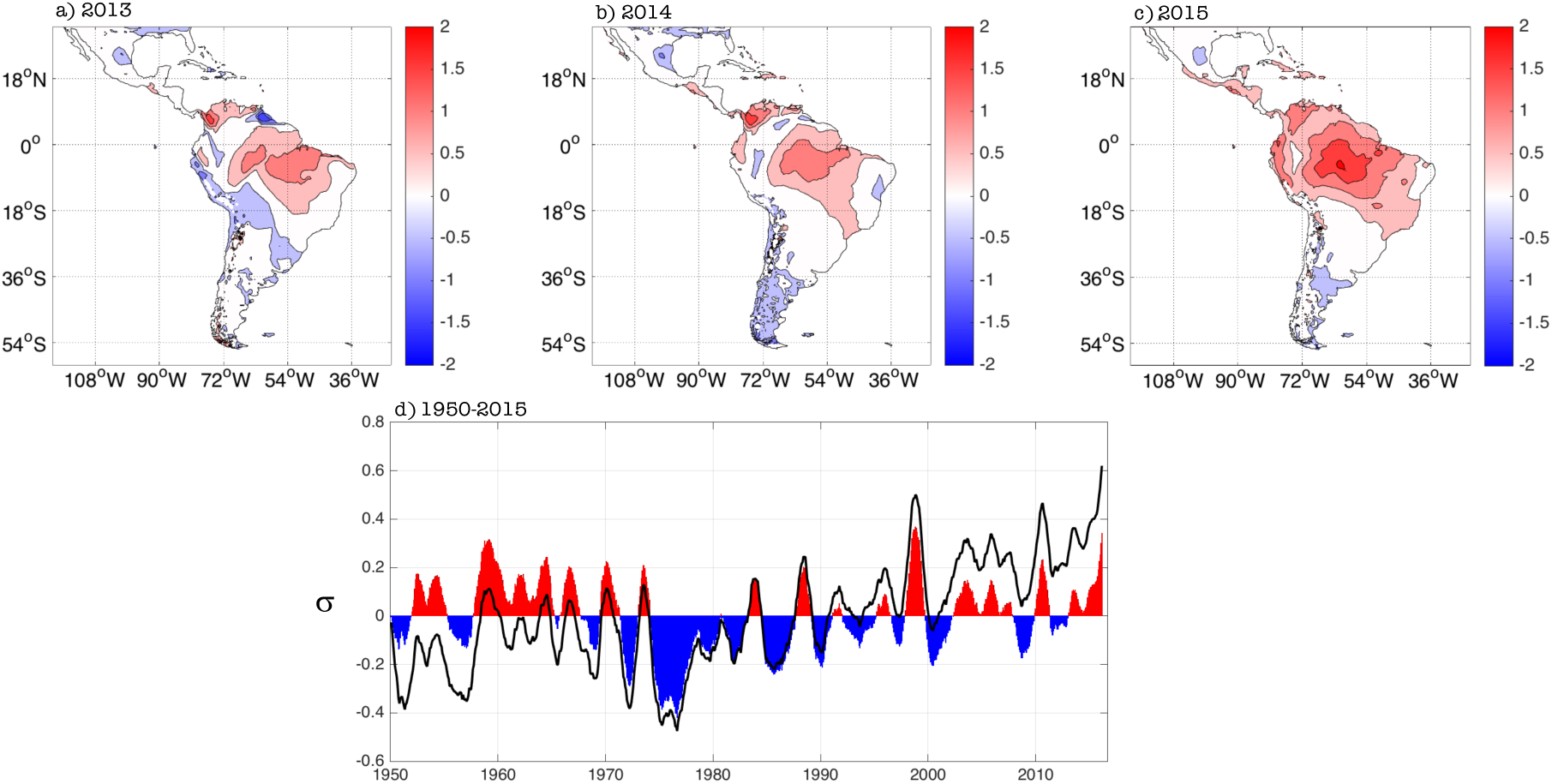
Spatial evolution of standardized ***R*_0_** yearly anomalies for (a) 2013, (b) 2014 and (c) 2015. (d) Average evolution of standardized ***R*_0_** anomalies (in units of standard deviations, ρ) for Latin America and the Caribbean (domain in panel (a)) for the 1950-2015 period. Black empty curve and filled curve show the raw and linearly detrended standardized anomalies, respectively. A 12-month running average filter was applied to both curves to better capture the inter-annual variability. There is no data over the oceans.

The neutral standardized anomalies in the Brazilian Nordeste (Northeast), one of the most impacted places in terms of the present ZIKV epidemic, are attributed to the buffering role of the Atlantic Ocean in controlling the local temperatures. Still, neutral standardized anomalies in Nordeste are associated with ***R*_0_** ranging between 3.5-5.5, which indicate a very high potential risk of transmission.

The high values of the 2015 standardized anomalies (Figure 3c) are also consistent with the observed behavior of other diseases like dengue; for example, the reported number of dengue cases for Ecuador in 2015 (42,667) was about 3 times larger than the average number of cases for 2011-2014 (14,467.5); for details see (39). Nonetheless, work of our team in Machala (coastal Ecuador) suggests that a high percentage of the 2015 dengue cases reported there are likely to be chikungunya cases. Even if that is the case, the model was able to capture enhanced conditions leading to a higher number of *Aedes*-borne diseases.

The evolution of the spatially-averaged ***R*_0_** standardized anomalies for Latin America and the Caribbean exhibits a clear trend between 1950 and 2015 (black curve in Figure 3d), as reported by (21), that we attribute to the persistent increase in temperatures observed in the region. Once the longer-term signals are filtered-out, the inter-annual component of the ***R*_0_** standardized anomalies (filled curved in Figure 3d), show a peak in 2015 that is the second-highest on record, after the one in 1998. This contrasts with the analysis performed by (21); overall Figure 3d is telling the same story as Figure 3 in (21), the main differences due to the use of a different dataset and mostly to the use of a 12-month running average in our case (see Section 2 above). Our interpretation is consistent with our previous study on the 2015 climate conditions (9): a superposition of long-term, decadal and inter-annual signals was responsible for the 2015 absolute maximum in the unfiltered time series (black curve in Figure 3d). Although most likely the 2015 El Niño had an important contribution, the maximum cannot be explained only by this inter-annual phenomenon.

## 4 Skill Assessment and DJF 2014-2015 Forecast

A new seasonal forecast system for potential risk of transmission of *Aedes*-borne diseases can be developed using the ento-epidemiological model discussed in the previous section and multi-model ensemble climate predictions at seasonal scale. For this purpose, we have selected the set of coupled global models participating in the North American Multi-Model Ensemble project (22). Although our focus is Latin America and the Caribbean, the same system can be used for other regions of the world, and a subset of the NMME models or a completely different seasonal climate forecast system can be used straightforwardly if that provides higher skill for the particular region of interest.

In brief, the system uses the monthly climate information from each one of the 116 (or 104, if the target period is between 2010 and 2015) realizations of the NMME models to compute the associated value of the basic reproduction number for each grid box in our geographical domain. Although the forecast horizon is typically 9 months after the initialization month, skill is normally higher for the first few seasons; to illustrate the approach here we focus on the first season starting immediately after the initialization month (e.g., JJA for forecasts initialized in May). After the multi-model ensemble and the seasonal average is computed, the output is corrected using a simple Principal Component Regression, which provided better results than other methods like Canonical Correlation Analysis or the use of the raw model output. For additional details, see Section 2.

The cross-validated analysis shows that there is relatively high skill (> 60%, as measured by the 2AFC metric) for ***R*_0_** for all the seasons considered in most of northern half of South America and several regions of Central and North America, and some Caribbean nations (Figure 4). Overall, the skill is higher in DJF and MAM (with Kendall’s t of 0.199 and 0.191, respectively), and minimum in JJA (0.123), SON being in the middle (0.146). These values of Kendall’s t are typical for rainfall predictions in the region, as can be seen in the Validation Maproom of the Latin American Observatory’s Datoteca (23,24,26): http://datoteca.ole2.org/maproom/Sala_de_Validacion/

**Figure 4.**
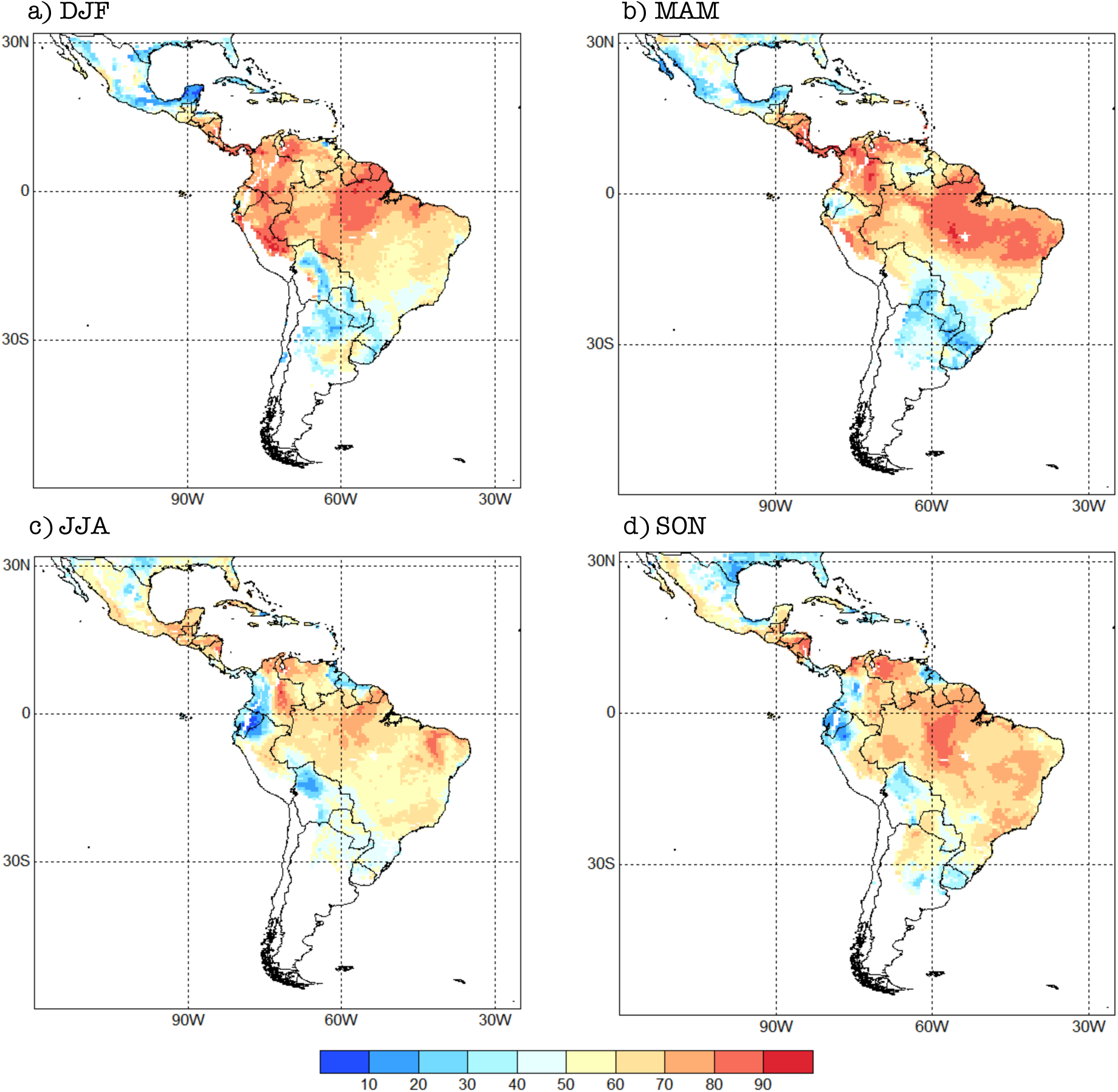
2AFC skill score for the seasonal forecast system for each one of the four seasons selected: (a) DJF, (b) MAM, (c) JJA and (d) SON. Units in %. The 2AFC score is an indication of how often the forecasts are correct; it also measures how well the system can distinguish between the above-normal, normal and below-normal categories.

Regionally speaking, skill is higher in Mexico in JJA, especially in the south (Figure 4). Central American countries exhibit high skill (above 70% for most of them) for DJF and MAM, with the lowest values (<50%) occurring in JJA and SON for Panama and Costa Rica. The western Caribbean tends to show higher skill during JJA, while the Central Caribbean and Lesser Antilles during MAM.

Northern South America shows relatively high skill (>70%) all year around, with some regions — like Ecuador, northern Peru, southwestern Colombia, northeastern Venezuela and northern Guyana— showing no skill during JJA and SON (Figure 4). The forecast system has in general low skill or no skill at all for southern South America, with some exceptions, e.g., the Bolivian Amazon in DJF, Paraguay and northern Argentina in SON, and northwestern Uruguay in DJF. Most of Brazil exhibits values of the 2AFC metric that are above 50% in all seasons, although southern Brazil has very low skill in MAM. In general, Chile and central and southern Argentina, do not show potential risk of transmission with this model, and thus those regions appear in white in our skill maps (Figure 4).

To illustrate an example of the bias-corrected probabilistic forecasts produced by our system, we now consider the season prior to the first ZIKV case reported in Brazil (May 2015 (40)): DJF 2014-2015. The probabilistic prediction indicates that there were mostly conditions for above-normal risk of transmission in eastern Brazil, which is similar to the observed conditions (Figure 5). Of course, no model is perfect and although, overall, there are similarities between the forecast and observed basic reproduction number for that season —especially in South America— there are also differences. Nonetheless, below-normal conditions were in general no forecast in the ZIKV hotspot places (in Brazil, for example), and as discussed above, the normal category the northern half of South America is already conducive to epidemic conditions. Hence, we claim that this particular forecast, even if not perfect, could have been useful for decision-makers at the time (November 2014), assuming that they already knew there was ZIKV in the region, which was of course not the case.

**Figure 5.**
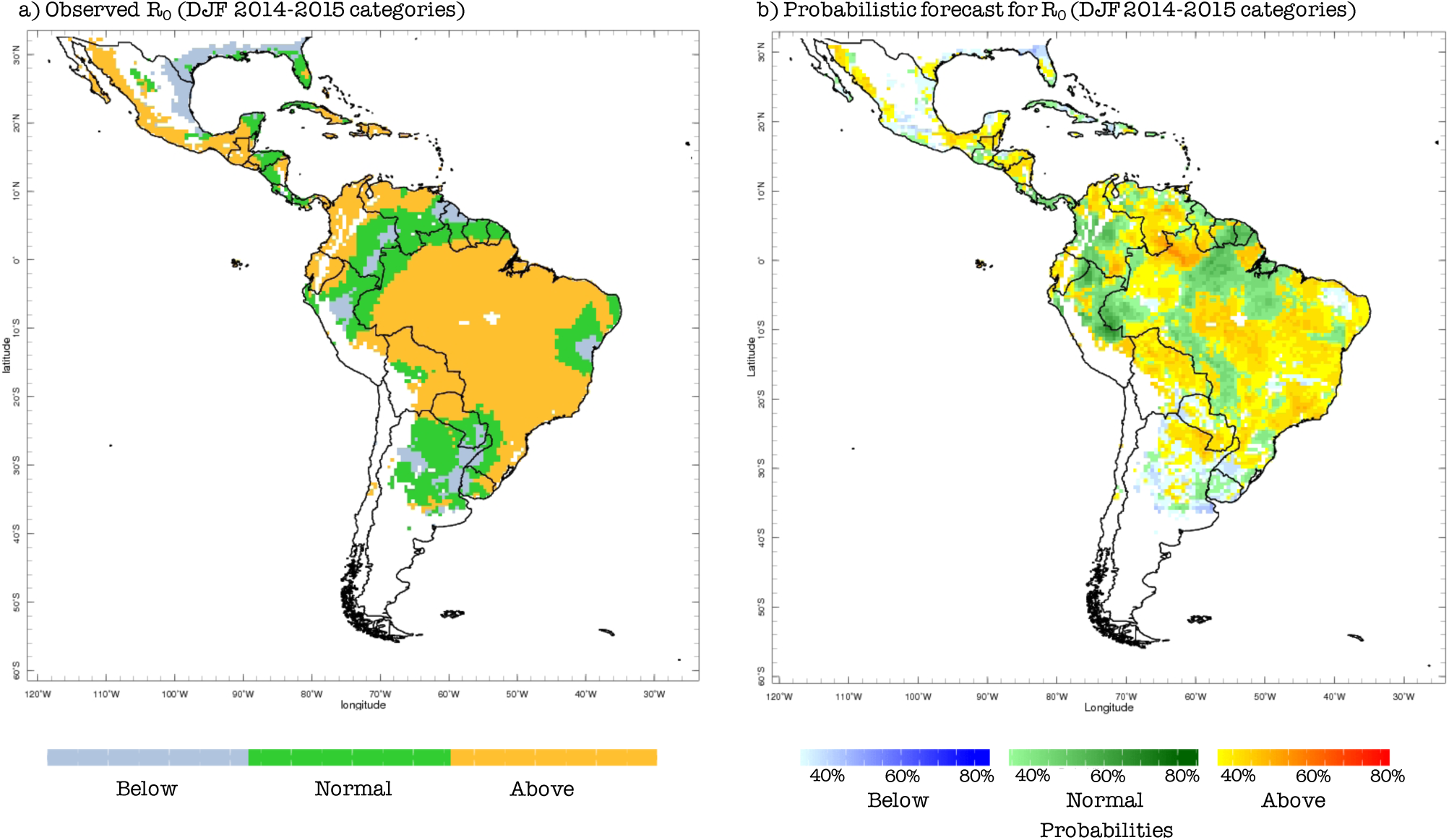
(a) Observed terciles (above normal, normal, below normal) for the basic reproduction number (***R*_0_**), computed using observed climate data for DJF 2014-2015 and the model presente in section 3. (b) Forecast probabilities (in %) for ***R*_0_** for the same DJF season, computed using predicted climate data, the vector model presented in section 3 and the probabilistic Principal Component Regression model described in section 2.

The previous example also illustrates why this tool can only be used as a guide for the local and international experts, as these diseases involve complex interactions beyond the presence or not of enhanced environmental (climatic) conditions suitable for the occurrence and transmission of *Aedes*-borne epidemics.

## 5 Concluding remarks

We have discussed the development and predictive skill of new probabilistic forecast system for above-normal, normal and below-normal environmental conditions associated with potential risk of transmission of diseases like ZIKV, DENV and CHKV. To the best of our knowledge this is the first seasonal forecast system of this type for Latin America and the Caribbean, although it is conceptually similar to a malaria forecast system developed for Africa years ago (18).

From the regional perspective, this forecast system has the potential to help the Pan-American Health Organization (PAHO), the World Health Organization (WHO) and other decision-makers to prepare more detailed epidemiological alerts and guides for zika’s surveillance and other arboviruses; to calculate different levels of population at risk and incidence rates for regional assessment, to prepare vector control guidelines for a more integrated management; to plan and support vector control resources an equipment; to organize and program activities and resource mobilization, as well as improve risk communication materials. One of the co-authors (PN) has already started to explore ways to take advantage of this forecast system at PAHO/WHO.

Our system is a first attempt to provide predictive tools for health practitioners and decision-makers interested in *Aedes*-borne diseases in Latin America and the Caribbean, and can be considered an additional step in the direction followed by previous research groups (21,34,36,41–44).

Indeed, forecasts of health events are designed to change human behavior. Nonetheless, as with the practice of medicine, there are ethical issues to consider. It is possible that there might be negative consequences from an epidemic risk forecast (i.e., incidence, or cases), even if the prediction is skillful. To illustrate this idea, consider that a forecast for ZIKV is provided to the community, indicating that there is above 80% probability of acquiring the disease in Rio de Janeiro during a certain season, but less than 10% probability of infection in Montevideo. People —some of whom could already be infected with ZIKV, or even with a different disease— might decide to travel to Montevideo instead of Rio de Janeiro because of that forecast, thus igniting or being part of a new focus of an epidemic there, that was not predicted and that is partially caused by the original prediction itself. This is an important caveat to be considered by the decision-makers. Another consideration is that a ZIKV forecast may have negative consequences for tourism, leading to livelihood impacts that may have negative health consequences.

There are a number of important limitations related to our forecast system. As indicated earlier in this paper, by itself this kind of system cannot forecast the occurrence and spread of new epidemics, but only partial conditions for that to happen. The model employed here only considers the effect of climatic conditions, through temperature and rainfall, on disease transmission via the vectors and viruses of interest. Direct human to human transmission via sexual intercourse and blood transfusion are outside the scope of this modelling approach. Also, the present version of the model cannot simulate co-infections or mixed states (e.g., a fraction of the population that is recovered from dengue but it is susceptible to zika).

One particular way in which the model needs to be improved involves how rainfall is considered. The present version of the model only uses rainfall in a rather simplistic way, without really considering its seasonal characteristics. There are examples in the scientific literature that could be used to improve the representation of rainfall in this type of model (see for example, (45–47)). In addition, there is room to consider a better set of realizations in the ensemble of simulations, varying the ento-epidemiological parameters of the model. This will be explored in the future.

## 7 Author Contributions

ÁM and MT established the concept of the study. ÁM obtained the data. All authors undertook the analysis and interpretation of results. ÁM, MT and AS drafted the manuscript. All authors critically reviewed and revised the manuscript and agreed the final submission.

## 9 Abbreviations

2AFC: Two-Alternative Forced Choice
CAMS: Climate Anomaly Monitoring System
CHIKV: chikungunya virus
CPT: Climate Predictability Tool
CRU: Climate Research Unit, at East Anglia University
DENV: dengue virus
DJF: December-January-February
JJA: June-July-August
IRI: International Research Institute for Climate and Society, at Columbia University
MAM: March-April-May
NMME: North American Multi-Model Ensemble
SON: September-October-November
PAHO: Pan-American Health Organization
PCR: Principal Component Regression
WHO: World Health Organization
ZIKV: zika virus

## 10 Acknowledgments

The authors are thankful to the editors of this special number of Frontiers for their kind invitation to contribute a paper, and to Ángel Adames, Hongai Zhang and two anonymous reviewers who helped improve the clarity of the original manuscript. This research used computational resources from the Latin American Observatory for Climate Events, in particular its Datoteca.

## Footnotes

1 A hindcast is a retrospective forecast, made using the same methodology of actual forecasts, but for a past period of time. They are usually produced to evaluate forecast skill.

2 We found a typo in Table 1 for the expression of **μ_1_** in (21): the correct value in the argument of the exponential function is 51.4 – 1.3 T.

